# An Integrative Framework for Detecting Structural Variations in Cancer Genomes

**DOI:** 10.1101/119651

**Authors:** Jesse R. Dixon, Jie Xu, Vishnu Dileep, Ye Zhan, Fan Song, Victoria T. Le, Galip Gürkan Yardimci, Abhijit Chakraborty, Darrin V. Bann, Yanli Wang, Royden Clark, Lijun Zhang, Hongbo Yang, Tingting Liu, Sriranga Iyyanki, Lin An, Christopher Pool, Takayo Sasaki, Juan Carlos Rivera Mulia, Hakan Ozadam, Bryan R. Lajoie, Rajinder Kaul, Michael Buckley, Kristen Lee, Morgan Diegel, Dubravka Pezic, Christina Ernst, Suzana Hadjur, Duncan T. Odom, John A. Stamatoyannopoulos, James R. Broach, Ross Hardison, Ferhat Ay, William Stafford Noble, Job Dekker, David M. Gilbert, Feng Yue

## Abstract

Structural variants can contribute to oncogenesis through a variety of mechanisms, yet, despite their importance, the identification of structural variants in cancer genomes remains challenging. Here, we present an integrative framework for comprehensively identifying structural variation in cancer genomes. For the first time, we apply next-generation optical mapping, high-throughput chromosome conformation capture (Hi-C), and whole genome sequencing to systematically detect SVs in a variety of cancer cells.

Using this approach, we identify and characterize structural variants in up to 29 commonly used normal and cancer cell lines. We find that each method has unique strengths in identifying different classes of structural variants and at different scales, suggesting that integrative approaches are likely the only way to comprehensively identify structural variants in the genome. Studying the impact of the structural variants in cancer cell lines, we identify widespread structural variation events affecting the functions of non-coding sequences in the genome, including the deletion of distal regulatory sequences, alteration of DNA replication timing, and the creation of novel 3D chromatin structural domains.

These results underscore the importance of comprehensive structural variant identification and indicate that non-coding structural variation may be an underappreciated mutational process in cancer genomes.

## Introduction

Structural variations (SVs), including inversions, deletions, duplications, and translocations, are a hallmark of most cancer genomes^1^. The discovery of recurrent SVs and their molecular consequences for gene organization and expression has greatly advanced our knowledge of oncogenesis. Numerous oncogenes have been identified as the product of recurrent translocations and have provided successful targets for drug therapies ^2-6^. Further, structural variations also provide clear diagnostic and prognostic information in the clinic ^7^. Despite their importance, comprehensively identifying structural variations in cancer genomes remains challenging, hindering our ability to better understand oncogenesis and to develop targeted treatments for cancer.

Several methods are currently used to identify structural variations in cancer genomes. G-band karyotyping has been the major method historically to detect gross structural anomalies in the genome, and it is routinely performed today clinically for certain malignancies such as leukemia ^8^. However, karyotyping is an inherently low resolution and low throughput method that cannot adequately characterize extensively rearranged genomes. Microarrays are another commonly used method for detecting gains and losses of genetic material ^9^, but they do not provide precise localization of rearrangements. Further, microarrays are inherently limited in detecting balanced rearrangements, such as inversions or balanced translocations. Finally, targeted approaches such as fluorescence *in situ* hybridization (FISH) and PCR are also used extensively in the clinic. However, these methods require *a priori* knowledge of the rearrangement events and hence are not suitable for *de novo* detection of structural variations.

Recently, high-throughput sequencing based methods have emerged as an attractive method for structural variant identification ^10, 11^. Targeted approaches such as exome sequencing and RNA sequencing provide cost-effective means of identifying gene fusion events ^12^, while whole genome sequencing (WGS) can provide information both on rearrangements as well as on gains and losses of genetic material ^13-16^. Despite their success, these high-throughput technologies are limited by their reliance on short sequence reads (usually less than 100 bp), which cannot be effectively mapped to the repetitive regions in the genome. More importantly, these techniques involve fragmenting the genome into approximately 500 base-pair fragments prior to sequencing, and as a result, much of the structural information in the genome is lost. Finally, in whole genome sequencing data structural rearrangements are represented only by the reads crossing the breakpoints greatly reducing the sensitivity of detecting of such events. Given the aforementioned limitations, it is imperative to develop alternative approaches for detecting structural variations that either use longer sequence reads or that retain long-range genomic structural information.

Here we propose an integrative framework to comprehensively detect SVs by using a combination of technologies, including WGS, next-generation optical mapping (BioNano Irys), and high throughput chromosome conformation capture (Hi-C). Although Irys and Hi-C have been previously used for genome assembly ^17-25^, this is the first time that WGS, optical mapping and Hi-C technology have been systematically used and integrated for SV detection in cancer genomes. Irys optical mapping works by first introducing single-strand cuts in DNA molecules with a sequence-specific nicking endonuclease, and then repairing the nick with fluorescently labeled nucleotides ^26^. Each DNA molecule is then straightened and electrophoresed through microfluidic nanochannels, through which DNA can migrate only when unfolded. Fluorescently labeled nicks are then imaged within the nanochannels. By aligning images of multiple DNA molecules at specific sites, this technology can generate high-throughput genomic maps for extremely long, single DNA molecules (∼200kb – 1Mb).

In addition to analyzing Irys optical mapping data, we develop novel algorithms to use Hi-C data to systematically identify structural variations genome-wide. Hi-C technology was initially invented to investigate genome-wide chromatin interactions ^27^ but has been recently adopted for other purposes, such as genome assembly ^19, 21^ and haplotype phasing. While the presence of structural variants has been observed in Hi-C datasets^19, 29-31^, we have developed and validated an approach to use Hi-C data to find structural variation in cancer genomes. We demonstrate that Hi-C can accurately detect structural variants in cancer genomes even with modest sequencing coverage (20-100 million reads or 1-5X coverage). We compiled a list of high confidence SVs in 8 human cancer cell lines by comparing the results from each of the three technologies, and we performed validation experiments on a subset of these variants. We observed that each method can detect distinct subsets of structural variations: Irys optical mapping and Hi-C excelled at detecting large and complex structural alterations, whereas high coverage WGS was adept at identifying smaller insertions, deletions, and rearrangements. Having obtained large-scale genomic structural information from Irys and Hi-C, we also investigated the consequences of these large-scale structural alterations in cancer genomes. We confirmed known oncogenic fusion transcripts such as BCR-ABL, and also discovered novel fusion events. We identified numerous instances of novel 3D chromatin structure alterations as a result of structural genome variation, such as the formation or dissolution of topologically associating domains (TADs), suggesting a novel role of structural variation in gene misregulation in oncogenesis.

## Results

### An integrated approach for structural variant detection

To comprehensively identify SVs in cancer genomes, we performed a combination of whole-genome sequencing, optical mapping, and Hi-C in 12 commonly used cancer cell lines representing different cancer types (Fig. 1a and Supplementary Table 1). First, we performed WGS in six cancer cell lines with an average coverage of 30X and obtained WGS in a seventh cell line from a previous study ^32^. We identified SVs using WGS data using an in-house pipeline mainly based on Manta software (Supplementary Table 2) ^33, 34^. Next, we performed optical mapping in eight cell lines with an average coverage of ∼100X, the most comprehensive optical mapping effort in cancer cells thus far. We used Irysview Refaligner to conduct *de novo* assembly and detect SVs for these cancer cell lines, and on average, identified ∼2,600 SVs in each cell line (Supplementary Tables 3). Lastly, we performed Hi-C experiments in 12 cancer cell lines and analyzed an additional 17 previously published Hi-C datasets (Supplementary Table 1)^29, 35-40^. We developed novel algorithms to identify potential re-arrangement events using Hi-C data (Supplementary Table 4). For this analysis, we focused on identifying inter-chromosomal translocations and intra-chromosomal re-arrangements greater than 1Mb apart, as our algorithm is less sensitive to SVs between regions within the same TAD (∼1 Mb) due to increased chromatin interactions ^41, 42^. Combining all methods, we detected thousands of insertions and deletions (>50bp), hundreds of inversions, and around 100 inter-and intra-chromosomal translocations in each genome (Table 1, Supplementary Table 5). We compiled a list of high-confidence SVs, which were predicted by at least two of the three methods (Supplementary Table 6). For example, Caki2 cells carry a translocation between chromosomes 2 and 3 that was detected by all three methods. This translocation was validated by observation of dramatic shifts in DNA replication timing profiles in this region (Fig. 1b). We obtained data from additional 8 cancer cell lines and 9 karyotypically normal diploid cells (Supplementary Table 1) and performed similar analysis. We visualized the high-confidence SVs as circular genome structural profiles^43^, which showed that the cancer genomes displayed many more rearrangement events compared with normal cells (Fig. 1c, Supplementary Fig. 1).

**Figure 1.**
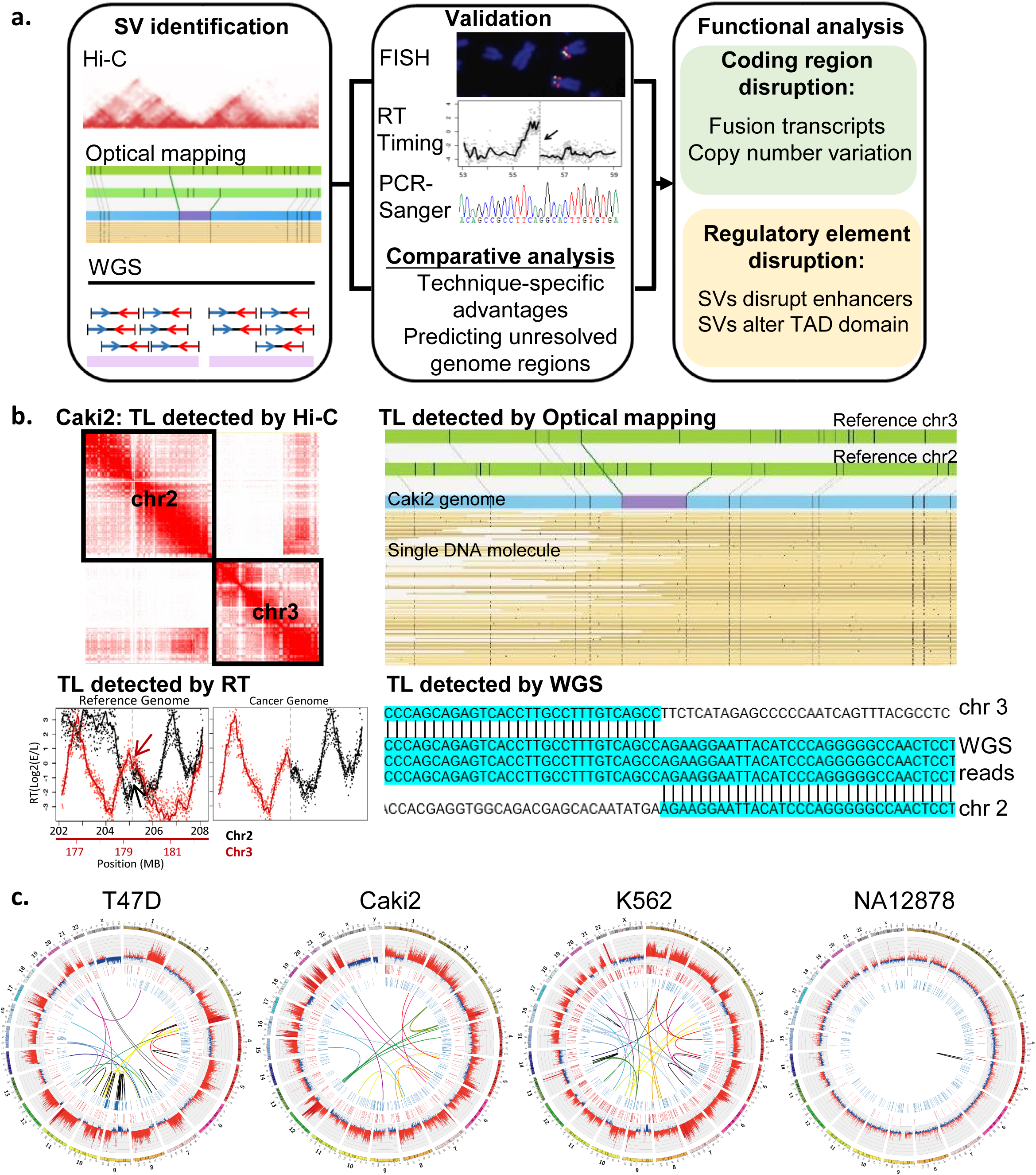
Overall strategy of SV detection in cancer genomes. **a.** The pipeline of SV detection, validation, and functional analysis. **b.** An example of SVs that are detected by all technologies (chr2: 205,125,031 and chr3: 179,412,688 in Caki2 cells). **c.** The cancer genome possess extensively more CNV and translocation in comparison with NA12878 (hg19). Tracks from outer to inner circles are chromosome scales, CNV, insertions, deletions and TLs. Outward red bars in CNV track indicate gain of copies (>2), and inward blue loss of copies (<2). CNV are profiled by Irys with 50,000 bp bin size. Insertion, deletion, and TLs are detected by at least two methods from WGS, Irys, and Hi-C.

**Table 1.** Number of SVs detected in T47D

| **No. of Events**<br>**(median size)** | **T47D** |  |  | **Non-redundant** |
| --- | --- | --- | --- | --- |
|  | **Irys** | **WGS** | **Hi-C** | **Total** |
| Deletion | 503 (4 Kb) | 4802 (84 bp) | - | 4922 |
| Insertion | 909 (2.5 Kb) | 2530 (108 bp) | - | 3175 |
| Inversion | 44 | 68 | - | 100 |
| Inter-chr translocation | 14 | 54 | 25 | 75 |
| Intra-chr translocation | 13 | 5 | 22 | 17 |

### Detection and Characterization of Translocations and Large Scale Re-arrangements

It has been suggested that strong inter-chromosomal interactions observed in Hi-C interaction matrices from cancer cell lines might be the result of SVs^19, 29-31^. While locus specific chromosome conformation capture has been used to identify whether individual genes are re-arranged in a given genome^44^, to our knowledge, no algorithm has been developed for systematic, unbiased, genome wide detection SVs from Hi-C data. We designed an iterative refinement algorithm to systematically detect translocations in cancer genomes using Hi-C data (details in method section, Supplementary Fig. 2). Fig. 2a shows an example of a potential translocation in Hi-C data. In normal cells, inter-chromosomal interactions are rare (Left panel in Fig. 2a). In Caki2 cancer cells, we observed strong “interchromosomal” interactions (Right panel in Fig. 2a), which are likely due to the fusion of chromosome 6 and chromosome 8. The challenge is to determine whether the increased signal is due to the translocated region forming chromatin interactions with its re-arranged partner, or the product of dynamic 3D genome organization. We have developed probabilistic models of Hi-C data to model “normal” features of 3D genome organization, including genomic distance between loci, TADs, A/B compartments, and the increased interactions between small chromosomes and between sub-telomeric regions. In the event of a re-arrangement, the two re-arranged regions are genetically fused, altering the linear distance between loci. This leads to local clusters of deviation from the expected interaction frequencies of the model. This signature can then be used for systematic identification of structural variation in any Hi-C dataset, including both inter-chromosomal translocations and intra-chromosomal re-arrangements (Fig. 2a,b).

We tested our method with a well characterized chronic myelogenous leukemia cell line (K562) and specifically used a limited number of sequencing reads (27 million read pairs and 1.54X coverage in replicate 1, generated as part of this study, and 22 million read pairs and 1.33X coverage in replicate 2, generated previously ^29^) to determine whether Hi-C can identify re-arrangements with limited sequencing depth. We started our analysis on large scale re-arrangements identified with a Hi-C bin size of 1Mb. We found 21 re-arrangements across the two replicates, of which 13 are known through prior karyotyping and the remaining 8 are novel re-arrangement events ^45^. The 8 novel re-arrangements were found in both replicates, suggesting that these are not a product of clonal evolution. More interestingly, several of them are novel complex re-arrangements: one event is between chromosome 16 and two different regions of chromosome 6 (Fig. 2c) and in another case, we observed a re-arrangement between chromosome 1, 6, 18, and 20 (Supplementary Fig. 3). We performed fluorescence in situ hybridization (FISH) experiments and validated a set of novel re-arrangement events. In total, 20 of the 21 predicted translocations using Hi-C data were validated by either FISH or previous karyotyping (Supplementary Table 7), suggesting that our algorithm can identify large-scale structural variation with high specificity. Next, to estimate the precise location of breakpoints, we iteratively applied the algorithm at increasingly smaller bin sizes to determine a subset of the re-arrangements with high resolution (Supplementary Fig. 4). For example, in K562 cells, we identified 4 of the 21 re-arrangements at 1kb resolution, all of which were further validated by PCR and Sanger sequencing (Supplementary Fig. 4).

To further evaluate the sensitivity of our approach, we evaluated its ability to detect the previously identified breakpoints on human chromosome 21 in Tc1 ES cells (Supplementary Fig. 5). Tc1 ES cells are a mouse ES cell line engineered to carry a copy of human chromosome 21^46^. In the process of establishing this cell line, human chromosome 21 was subject to gamma irradiation ^46^, leading to massive genomic re-arrangements, a subset of which have been previously identified using PCR and Sanger sequencing ^47^. We generated high coverage Hi-C data in Tc1 cells and performed our algorithm (Supplementary Fig. 5 a). By sub-sampling the data, we evaluated the sensitivity of our algorithm at various sequencing depths. The sensitivity ranges between 40%-90% depending on the sequencing depth and method used to call overlap, and appears to plateau when using 100 million sequencing read pairs or more (Supplementary Fig. 5b). In addition, we observe high internal consistency of breakpoints calls when there is at least 50 million reads (Supplementary Fig. 5c,d). This result suggests that our method requires only modest sequencing depths to achieve high sensitivity and saturation of breakpoint calls, and that we can achieve decent sensitivity with as little as 5-10 million reads. By examining the “missed” breakpoints, we observe that Hi-C may call breakpoints in identical regions as identified by WGS, but identifies a different strand as part of the breakpoint (Supplementary Fig. 5e,f). This suggests that although Hi-C does not always precisely resolve the strandedness of a subset of breakpoints, it may retain more information regarding the large-scale structure of the re-arrangement. Having demonstrated the sensitivity and specificity of our approach, we expanded our Hi-C analysis to 19 additional cancer cell lines and 9 karyotypically normal lines (Fig. 2d). We observed on average 28 re-arrangements in cancer cells and virtually no such events in normal cells. The rare instances of re-arrangements in normal cells all occur immediately adjacent to centromeres and therefore potentially represent anomalous or polymorphic assembly differences.

To systematically characterize the translocations in these cancer cells, we compared the translocations predicted by Hi-C, optical mapping, and WGS, compiling a list of high-confidence SVs in seven cancer cell lines where we performed all three experiments. Compared with previously known karyotypes in each lineage, the majority of the translocations detected in this study are novel Fig. 2e). We selected eight novel translocations of T47D for further validation, all of which were confirmed by PCR experiments (Supplementary Table 7).

We observed that 25% of these rearrangements were identified by more than one method (Fig. 2f). There is a large overlap between the translocations detected by Irys and Hi-C: 64% (9 out of 14) of inter-chromosomal translocations and 83% (10 out of 12) of intra-chromosomal re-arrangements detected by optical mapping in T47D cells are predicted by Hi-C (Fig. 2f, Supplementary Fig. 6). The overlap between Irys with WGS is much smaller: 33% (5 out of 14) of Irys-detected inter-chromosomal translocations are captured by WGS (Fig. 2f). The difference might be due to two reasons. First, the sizes of the rearrangements identified by WGS are smaller than those identified by optical mapping: 49% of WGS-specific translocations are less than 1kb. Since Hi-C and optical mapping both rely on either restriction enzymes or nickases that recognize target motifs with a spacing greater than 1kb (Hi-C) and 10Kb (optical mapping), small SVs are unlikely to be captured by these methods. On the other hand, both Hi-C and optical mapping can identify rearrangements in repetitive regions of the genome, which are usually missed by WGS. For example, one of the rearrangements in K562 cells is located near a centromere of chromosome 20, which contains many centromeric repeats and therefore is unmappable with WGS. However, Hi-C was able to leverage reads from nearby, mappable portions of the genome to detect the centromere-proximal breakpoint (Supplementary Fig. 3a), which we subsequently confirmed by FISH (Supplementary Fig. 3b). Likewise, since optical mapping operates on DNA fragments on the order of 100-200kb in size, minimizing the influence of short, repetitive sequences, it can discover many breakpoints in or near unmappable regions of the genome (Supplementary Fig. 6h). Finally, we observed that both Hi-C and Irys are particularly powerful at detecting complex translocation events, as illustrated in Supplementary Fig. 3. In summary, these results illustrate that each method has unique strengths and weaknesses, and should be applied accordingly depending on the study goals. Whenever possible, an integrative approach of different methods is essential to gain a more complete knowledge of structural variation in cancer genomes.

### Validation of breakpoints using replication timing

Eukaryotic genomes replicate via the synchronous firing of clusters of origins, which together produce multi-replicon domains each of which completes replication in a short burst during S-phase. Genome-wide profiling of replication timing reveals that these domains can be replicated at different times during S phase, with adjacent earlier and later replicating domains punctuated by regions of replication timing transition ^48, 49^. Consequently, translocations that fuse together domains of early and late replication can result in the earlier replication of the late replicating domain and/or delayed replication of the early replicating domain ^50, 51^. When mapped to the reference genome, these changes appear as abrupt shifts in replication timing profiles that have the potential to validate breakpoints (Fig. 2g). Our Hi-C pipeline identified 245 translocations in 9 cell lines in which replication timing is available. Out of these, 50 translocations were associated with an abrupt shift in replication timing. Since an abrupt shift is only expected for translocation between domains that replicate at different times, we aimed to classify the translocations based on the replication timing of the loci. However, the lack of a control cell line that represents the pre-translocation replication timing of the loci confounds this classification. To circumvent this problem, we classified the genome into regions that are constitutively early replicating (CE), constitutively late replicating (CL) and regions that switch replication timing during development (“switching regions”), using 35 replication timing profiles of non-cancerous cell types spanning all three embryonic lineages (Methods, Supplementary Fig. 7a) ^52, 53^. Among the 245 translocations detected by Hi-C, 17 were CE to CL fusions and 30 were CE to CE or CL to CL fusions. As expected, an abrupt shift in timing was identified in CE to CL with a much higher frequency (∼59%) than in CE to CE or CL to CL fusions (∼13%)(Supplementary Fig. 7 b, c). The majority (∼76%) of the translocations that could be classified as CE, CL or S involved a switching region making it difficult to confidently predict their replication timing before the translocation event.

**Figure 2.**
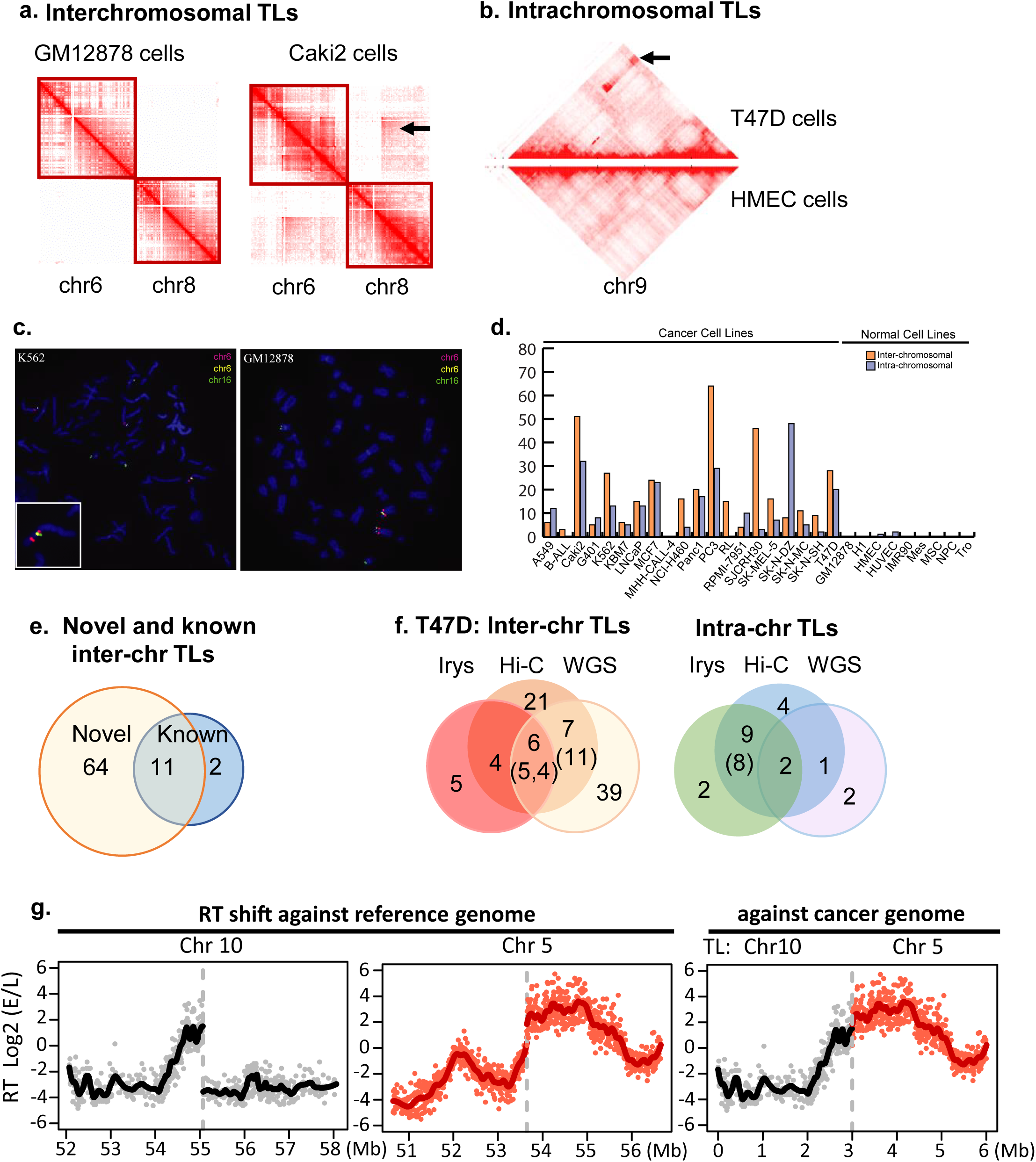
Detection and characterization of SVs in cancer genomes. **a-b.** Interchromosomal and intra-chromosomal TLs detected by using Hi-C data (←). **c**. Validating a complex TL (chr6-chr6-chr16) by FISH in K562 cells. **d.** Inter-chromosomal and intrachromosomal TLs detected by Hi-C in 20 cancer genomes and 9 normal genomes. **e.** Overlap between known and novel TLs in T47D cells detected in this study. **f.** Comparison of interchromosomal and intra-chromosomal TLs detected by three methods. A large portion of WGS-specific TLs are smaller than 10Kb. **g.** The impact of TLs on replication timing. RT profiles of chr5 and chr10 of SKNMC, when plotted to the reference genome, show abrupt shifts at the TL breakpoints (←, left panels), and they are smoothly connected due to their juxtaposition in the cancer genome (right panel).

### Detection and Characterization of Copy Number Variations

We also identified numerous gains or losses of genetic material by optical mapping and WGS. First, we observed that optical mapping detects significantly fewer deletions than WGS (504 vs 4804, Fig. 3a, Supplementary Fig. 8a, b), but the sizes of Irys-detected deletions are larger (median size 7 kb vs 100 bp, Fig. 3b). Thus, 94% (4425/4723) of WGS-detected deletions cannot be predicted by Irys, because optical mapping relies on nickases that recognize specific sequence motifs and cannot detect small indels. However, 35% of the Irys-detected deletions are not captured by WGS, because optical mapping retains long-range contiguity of large DNA fragments (>100kb) and can identify larger deletions that are frequently missed by WGS. We tested a subset of Iry-specific deletions and 87.5% (14 of 16) were validated by PCR (Supplementary Table 7). In addition, optical mapping can identify deletions in repetitive regions that WGS misses, as shown in Fig. 3c. Furthermore, the deletions identified by Irys are enriched for repeat elements (compared to the genome background), whereas those identified by WGS show little or no enrichment for repeats (Fig. 3d). Hi-C data was not used for CNV analysis, as it does not provide any obvious additional information than WGS for this purpose.

To further investigate whether the deletions are specific to cancer genomes, we compared the deletions detected in this study with the Database of Genomic Variants (DGV), which contains known SVs identified by previous studies in healthy individuals, including the three phases of the 1000 Genomes Project. The majority (85%) of deletions identified in cancer cell lines have been previously identified in the DGV (Fig. 3e), suggesting that many of the deletions represent polymorphisms in the population. However, we observed that cancer cells suffer more loss of genetic material (Supplementary Fig. 8c), and exhibit a greater number of rare and novel re-arrangements in comparison with GM12878 (40% vs. 10%), suggesting that a portion of the SVs represent somatic mutation events in cancer cells. The previously identified polymorphic deletions from the healthy population are enriched for repetitive elements (70% vs 50% genomic background) and depleted of functional elements such as enhancers (10% vs 20% genomic background) and exons (2.5% vs 1% genomic background) (Supplementary Fig. 8d-f). In contrast, the novel deletions are not enriched in repeats or depleted of enhancers or exons. Instead, they are enriched in COSMIC tumor related genes (Fig. 3e)^54^, suggesting that a subset of the deletions are potentially pathogenic.

We also compared the ability of WGS and optical mapping to detect global patterns of copy number variations. Individual fragments generated by optical mapping can be used to detect changes in depth of coverage over given regions, similar to WGS or microarray-based approaches for CNVs detections. By analyzing chromosome-wide patterns of copy number changes, we detect strong concordance between the results of optical mapping and WGS (Supplementary Fig. 9).

### Better Estimation of Gap Regions in Human Reference Genome

Interestingly, we discovered that optical mapping can help refine genome assemblies, especially with respect to estimating the size of gap regions. First, we noticed that the number of Irys-detected structural variants differs substantially when we compared the results to different version of the reference genomes (hg19 vs. GRCh38). Further investigation shows that many predicted “deletions” in hg19 by optical mapping consist of gaps in the reference genome that have been corrected in the GRCh38 build. The corrected size in GRCh38 is very similar to the prediction by optical mapping based on deletion detection (Supplementary Table 8), suggesting the power of optical mapping in accurately estimating the size of gap regions. For example, when we used hg19 as the reference genome, optical mapping in 10 cell types (4 normal primary cells and 6 cancer cell lines) recurrently predicted a 143Kb “deletion” within genomic loci chr1: 3,845,268-3,995,268. This region, however, is annotated as a 150Kb gap. Therefore, we predicted that the real gap size in the human reference genome should be 6.68Kb. When we checked the GRCH38 reference genome, we found that the size of this gap has been corrected to 6.51Kb, which is strikingly similar to the value we predicted.

However, we noticed that there remain several such “deletions” over gap regions even in the GRCh38 build that are not consistent with our findings with optical mapping, indicating that these gap sizes may still be in error or that there may be individual heterogeneity in gap sizes over these regions. The improved gap size estimation for GrRch38 is provided in Supplementary Table 9.

### Fusion transcript and gene dosage in cancer cancers

We investigated the functional consequences of these genetic alterations identified in cancer cell lines. First, we examined gene fusions due to genomic rearrangement, with a goal to both confirm known events and identify novel ones in these cancer cells. We analyzed RNA-Seq data of 11 cancer cell lines, including K562, and investigated whether we can detect fused gene transcript that are consistent with the genomic rearrangements identified in this study. We detected numerous RNA-Seq read pairs whose two ends are mapped to different chromosomes, crossing the identified translocation breakpoints (Supplementary Table 10). We confirmed some known oncogene transcripts, such as the *BCR-ABL* gene fusion. Importantly, we discovered many novel fusion transcripts involving bona fide oncogenes, such as EVI1-CFAP70 in T47D cells, whose expression was increased more than 10 fold compared with the un-translocated genes. How these novel gene fusions events contribute to the oncogenic potential remains to be further investigated.

Copy number alterations also represent a well-defined class of genetic variation in cancer. Prior studies have shown the presence of recurrently amplified and deleted genes in diverse cancer types ^9^. Examining the deleted regions that we identify by WGS and Irys, we observed that deleted regions are enriched for Gene Ontology annotations related to tissue-specific and cancer type-specific genes and pathways (Fig. 3f, g). Further, by comparing with the recent findings from WGS data of 560 breast cancer patients ^16^, we observed that 8 out of the top 10 frequently mutated oncogenes in breast cancer patients were also amplified in T47D cancer cells, and tumor suppressor genes such as *ATRX* and *CDKN1B* displayed loss of copies (Fig. 4a), suggesting that T47D cells reflect the CNV landscape in breast cancer and our method can accurately capture these variations. We further compared the RNA-Seq data in T47D cells with those from human mammary epithelial cells (HMEC), confirming that loss-of-heterozygosity (LOH) and homozygous deletions in T47D cells indeed lead to significantly reduced gene expression correlated to the number of lost copies (Fig. 4b). For example, one 18Mb deletion in T47D results in LOH of over 400 genes, and decreased transcription of the majority of this set of genes (Supplementary Fig. 10a,b). We found deletions in exonic regions of a total of 25 COSMIC tumor-related genes, and the majority (76%) showed decreased transcription (Supplementary Fig. 10c). We made similar observations when comparing transcriptomes in other cancer cells (Supplementary Fig. 10d). These results suggest that our integrated method can accurately capture changes in gene dosage in cancer genomes. As we extended the CNV analysis onto 7 cancer cell lines, we noticed widespread amplification of known oncogenes (such as *MYC*) and loss of cell cycle checkpoint genes (such as CDKN2A/B, Supplementary Fig. 11). In addition, we found over 100 genes that are extensively amplified or deleted in cancer cells but were not reported in COSMIC, suggesting their potential roles in cancer (Supplementary Fig. 12).

**Figure 3.**
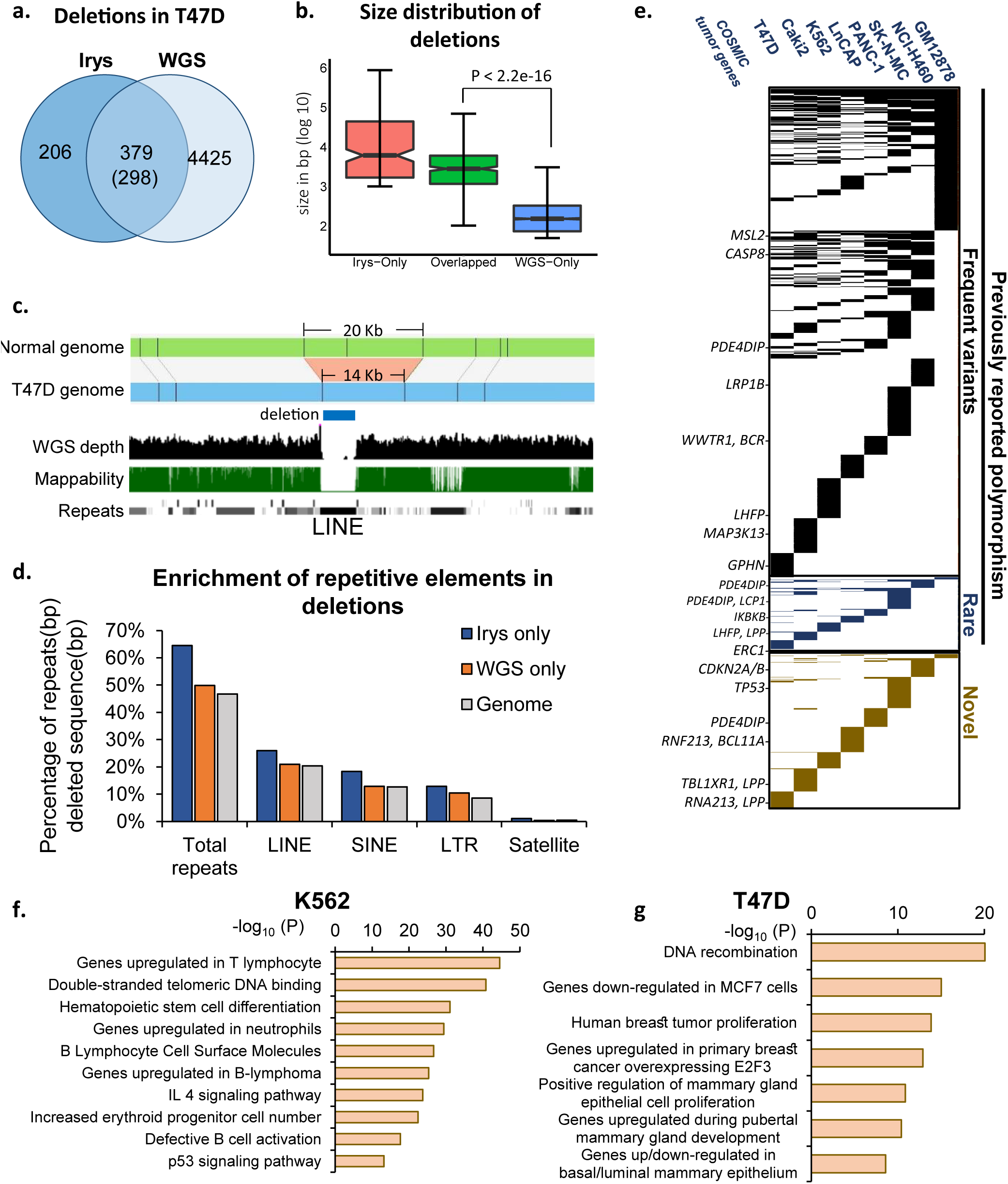
Comparative analysis and characterization of deletions in cancer genomes. **a.** Overlap of deletions detected by optical mapping and WGS. **b.** Size distribution of deletions detected by optical mapping and WGS. **c.** Optical mapping detects a 6Kb deletion within chr15:55,215,578-55,235,682 that is missed by WGS. The deletion is located inside an unmappable region and overlaps with a LINE element. **d.** Deletions detected by Irys are enriched of repetitive elements in comparison with genomic background and deletions detected by WGS. **e.** Over 85% of deletions (detected by both WGS and Irys) from cancer and normal cells represent polymorphism, including frequent variants (reported 7∼11813 times from population of healthy individuals) and rare variants (reported 1∼6 times). Rare and novel unreported variants account for less than 10% of deletions in normal genome, but account for 40% of deletions in cancer genomes. Cancer-specific, rare, and unreported variants are enriched of COSMIC cancer-related genes. **f-g**. Deletions detected in K562 and T47D are located near tissue-specific and cancer-specific genes.

**Figure 4.**
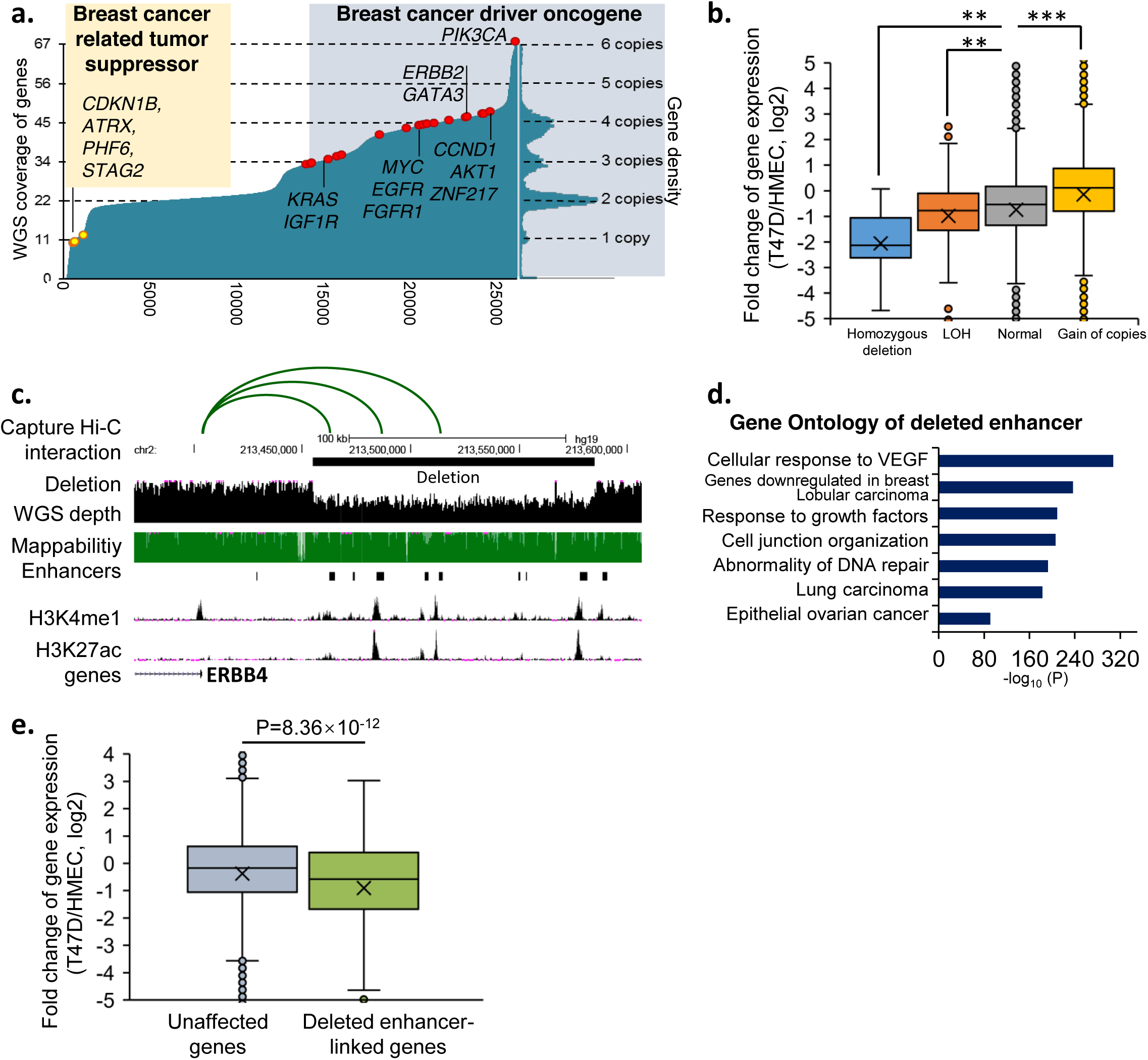
The impact of CNV on transcription. **a.** The top 93 breast-cancer related genes show extensive amplification of oncogene and loss of tumor suppressor genes. Shown in the figure are all RefSeq genes sorted by copy number. **b.** Genes with homozygous deletion and LOH in T47D show reduced expression (p=0.009, p=0.003), and genes with gain of copies increased expression (p=4×10^−79^), relative to the expression in HMEC cells. **c.** A 130Kb deletion in T47D contains a cluster of distal regulatory elements, three of which are linked with ERBB4 gene by Capture Hi-C. **d.** The deleted enhancers are located near genes important for pathways related to breast cancer. **e.** Genes with deleted enhancers show reduced expression levels. We only used gene without exon deletion or copy number loss in this analysis. Then we put genes into two groups: 534 genes with loss of at least one linked enhancers indicated by Capture Hi-C; 10677 without deletion of linked enhancer.

### Disruption of non-coding elements in cancer genomes

We are also interested in whether structural variants can affect non-coding regulatory elements and whether such alterations play a role in oncogenesis. For this analysis, we focused on comparing the enhancer landscape in T47D breast cancer cells and HMEC cells. We downloaded histone modification data from both cell types from the ENCODE Consortium and then predicted candidate enhancers based on H3K27ac signals. By comparing the enhancer annotations in HMEC and the deleted regions in T47D, we identified potential deleted enhancers in T47D cancer cells (Supplementary Table 11). We show an example in Fig. 4c, where multiple strong candidate enhancers present in HMEC cells are in a region deleted in T47D cells. These candidate enhancers are located upstream of gene *ERBB4*, which has been shown to play a crucial role in breast cancer. These candidate enhancers are linked to the promoter for *ERBB4* in recently published capture HiC data^55^ (Fig. 4c), lending support to our hypothesis that deletion of this region could play a role in breast cancer via effects on *ERBB4* expression. To investigate whether candidate enhancer elements deleted in T47D cells are broadly associated with cell growth control, specifically whether they affect any known signaling pathways, we performed Gene Ontology analysis with the GREAT tool. Strikingly, we found that these deleted enhancers of T47D are located near genes important for cellular response to VEGF, genes down-regulated in breast cancer, and genes involved in abnormality of DNA repair (Fig. 4d). Furthermore, we observed that genes linked to these deleted enhancers by capture Hi-C in HMEC cells show a reduced level of expression in T47D breast cancer cells (Fig. 4f). Overall, these results suggest that deletions in cancer genomes may frequently remove enhancers and thereby contribute to oncogenesis. Whether these enhancer deletions represent recurrent alterations to cancer genomes remains to be further investigated in patient samples and validated by additional functional experiments.

### The impact of structural variation on 3D genome organization

Our previous work in karyotypically normal cells and tissues has suggested that topologically associating domains (TADs) are fundamental features of 3D genome structure that are conserved in diverse cell types and species. Several recent reports have shown that genetic mutations can disrupt TADs and create “neo-TADs”^56, 57^ that in turn can lead to misregulated gene expression in developmental disorders ^56, 57^. Further, recent reports have also indicated that deletion of CTCF binding sites can disrupt local looping events leading to mis-regulation of nearby oncogenes ^58^. However, how SV contributes to changed 3D genome organization in cancer remains elusive.

Having identified structural variants in 20 cancer cell lines with Hi-C data, we systematically investigated the consequences of structural variation on TAD structure in cancer genomes. We observed that neo-TADs are formed as the result of large-scale genomic re-arrangements in cancer cells (Fig. 5a,b). For example, we identified a translocation from chr7 to chr8 in SK-N-SH cells (Fig. 5a). The breakpoint region on chromosome 8 occurred roughly 300kb downstream from the MYC/c-myc gene. SK-N-SH cells are a neuroblastoma cell line, and whereas most neuroblastomas express high levels of n-myc, SK-N-SH cells are a rare sub-type that express high levels of c-myc but not n-myc ^59^.

In examining the 3D genome structure near this re-arrangement, we observed extensive interactions in the vicinity of the re-arrangement on chr7 and chr8 that appear to form a neo-TAD. This neo-TAD extends ∼300kb upstream of the breakpoint to precisely the location of the MYC/c-myc gene, indicating that this translocation may form a neo-TAD which includes MYC as well as regions over 1Mb across the breakpoint junction. We generalized this observation by averaging the interaction signal across all translocation breakpoints in all cell lines (Fig. 5c). We observed that on average, the nearest “normal” TAD boundaries appear to be fused together on each side of the breakpoint creating neo-TADs (Fig. 5c). We analyzed gene expression profiles of 16 cancer cell lines for which RNA-seq data is available. We observe more variable expression patterns of genes when they reside within TADs containing a re-arrangement in the same cell type (Fig. 5d). These results suggest that re-arrangements within TADs may lead to aberrant expression patterns of genes within the TAD. These results indicate that structural variations in cancers can re-wire TAD structure to create novel domains in cancer genomes and potentially leading to altered regulatory environments within the domain (Fig. 5e). Determining whether any individual neo-TAD represents a re-current alteration in a given cancer cell type, or how neo-TADs may ultimately contribute to oncogenesis, remains to be elucidated. However, our analysis suggests that creation of neo-TADs is a common consequence of re-arrangements in cancer genomes.

**Figure 5.**
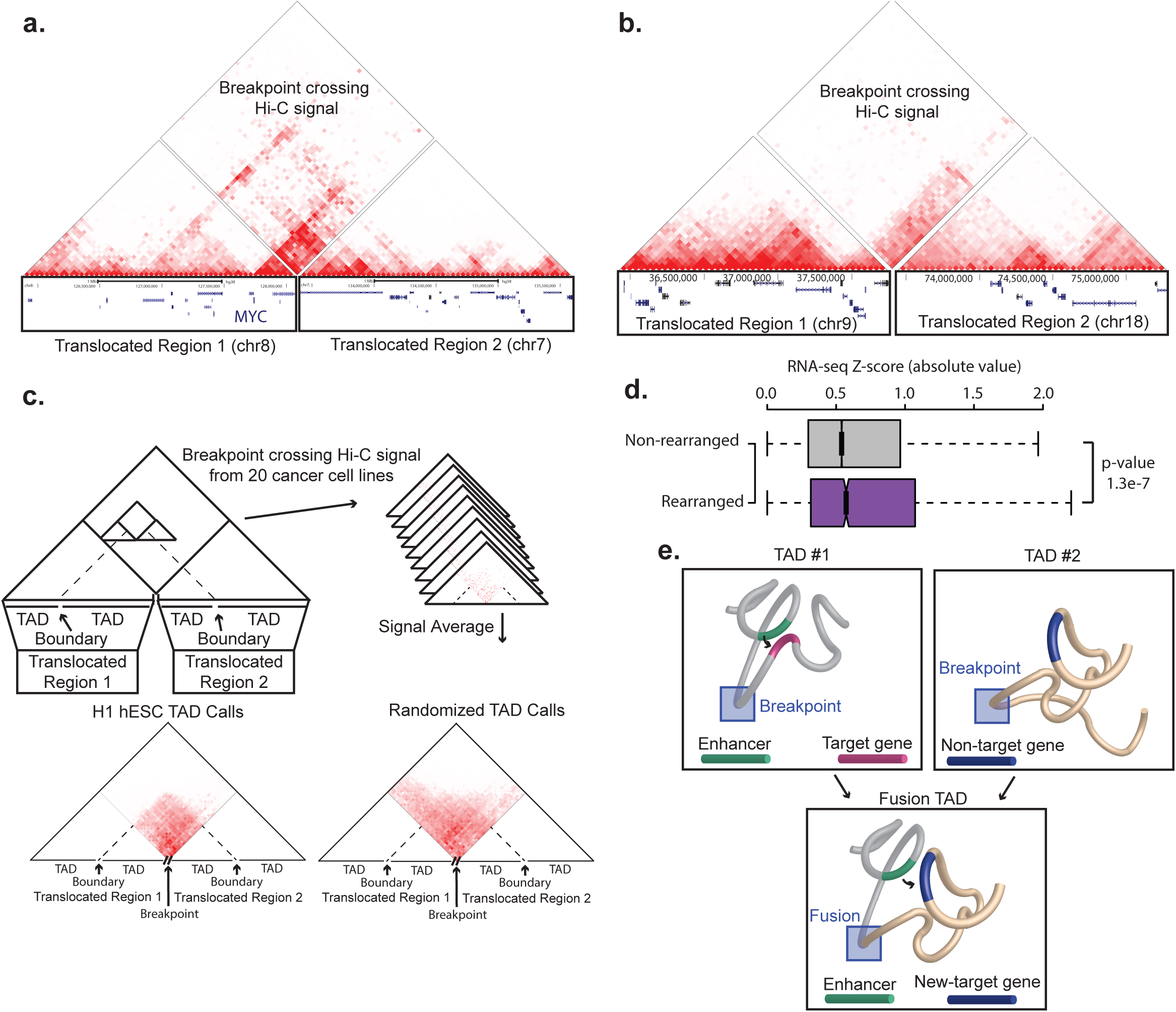
Re-arrangements and TAD fusions. **a.** Hi-C signal in near a chromosome 7:8 translocation in SK-N-SH cells. The left hand region of the genome browser track shows the re-arranged region on chromosome 8, with the location of the MYC gene marked, while the right hand region shows the area on chromosome 7. The tracks are arranged relative to each other to represent the predicted genetic structure of the re-arranged allele, with the breakpoint fusion site in the middle. Above the tracks is the Hi-C signal. The triangle heat maps on the left and right show intra-chromosomal interactions near the re-arranged regions. The diamond shaped heat map shows the Hi-C signal that crosses the breakpoint. The breakpoint crossing Hi-C signal shows the presence of a new TAD formation between the re-arranged regions. **b.** Similar to panel a, showing a TAD fusion after a translocation between chromosomes 9 and 18 in Pancl cells. **c.** Aggregate analysis of TAD fusions. The breakpoint crossing Hi-C signal across all breakpoints in all 20 cell lines was averaged and centered on the interacting bins between the nearest TAD boundaries (left) or randomized TAD boundaries (right). The average signal from the true TAD boundaries shows a marked enrichment within the regions demarcated by the closest TAD boundary, suggesting that TAD fusions are a common result of re-arrangements in cancer genomes. This demarcation is lost when using randomized TAD boundaries. **d.** Boxplot showing the distribution of gene-wise Z-scores of genes located in TADs containing re-arrangements (Rearranged) or not (Non-rearranged). The log2 of gene RPKM expression values was converted to a gene-wise Z-score for each gene across all 16 cancer cell types with Hi-C and RNA-seq data. The boxplot shows the absolute value of these Z-scores. Gene located in TADs containing rearrangements tend to have higher absolute value of their Z-scores, indicating more variable expression patterns. **e.** Model for neo-TAD formation. Native TAD structures are rearranged as the result of breaks and fusions, resulting in the juxtaposition of regulatory sequences from one domain and genes from another, potentially altering the regulatory landscape of cancer genomes.

## Discussion

Comprehensively detecting structural variations in cancer genomes remains a challenge for geneticists and cancer biologists. Here, we developed an integrative approach that employs a combination of WGS, optical mapping, and Hi-C to detect structural variations. We applied the procedure to three cancer genomes and validated a selected subset of them by PCR and FISH. No single method comprehensively identifies all structural variants, and each approach has its own strengths and weaknesses. Hi-C is extremely sensitive for detecting inter-chromosomal rearrangements and can readily detect rearrangements greater than 1Mb in size on the same chromosome. Furthermore, the algorithm we developed in this work does not require deep sequencing of Hi-C, successfully detecting rearrangements with little more than ∼1X coverage of the genome. However, our algorithm currently has limited power in detecting alterations less than 1Mb in size. On the other hand, optical mapping can readily detect both intra-chromosomal and inter-chromosomal alterations, including rearrangements less than 1Mb in size. Furthermore, optical mapping can be used to detect CNVs, similar to what is commonly done with WGS-or microarray-based approaches. However, optical mapping cannot identify small deletions and insertions (< 1kb). Finally, WGS can detect small deletions and insertions, and has higher resolution than Hi-C or optical mapping. However, WGS is less successful with detection of SVs in poorly mappable regions of the genome or complex structural variants.

In examining regions affected by structural variants identified in this study, we observed well-characterized functional consequences, such as the creation of gene fusions and changes in gene dosage in cancer genomes. In addition, we detected extensive deletions of distal enhancer elements. These deletions are enriched for proximity to genes known to be mutated in cancer and important for pathways in cancer biology, including DNA repair and signal transduction. To what extent such distal non-coding mutations are re-current in cancer genomes remains unclear, but this represents an important largely unexplored aspect of cancer genomics. Lastly, by analyzing the 3D genome structure surrounding the structural variants, we observed the creation of new TADs as a result of genomic rearrangements in cancer genomes. TADs appear to be an invariant organizational principle of metazoan genomes, and alterations that disrupt TAD structure have already been shown to underlie certain rare disorders of limb development. There is ample evidence that the juxtaposition of active regulatory sequences to known oncogenes can contribute to tumorigenesis. These results indicate that at least part of this effect may result from the creation of novel structural domains in cancer genomes.

## Acknowledgements

This work was supported by NIH grants U01CA200060 (F.Y.), R24DK106766 (R.C.H. and F.Y.), GM083337 (D.M.G.), GM085354 (D.M.G.), DK107965 (D.M.G.), U41HG007000 (W.S.N.), HG003143 and DK107980 (J. Dekker), DP5OD023071 (J. Dixon). This work was also supported by European Research Council (D.T.O., C.E. #615584), Cancer Research UK (D.T.O., C.E. #20412 & 22398), Wellcome Trust (D.T.O, C.E., S.H. #84459, #106985). J. Dekker is an investigator of the Howard Hughes Medical Institute. J. Dixon is also supported by the Leona M. and Harry B. Helmsley Charitable Trust grant #2017-PG-MED001. F.A. was supported by Institute Leadership Funds from La Jolla Institute for Allergy and Immunology. F.Y. is also supported by Leukemia Research Foundation and Penn State Clinical and Translational Science Institute. We thank B. Zhou and A.E. Urban (Stanford University) for sharing the SV profiles in K562 cells. We thank the ENCODE Data Coordination Center for helping with Hi-C and replication time data deposition.

## Author Contributions

J.R.D., J.X., and V.D. led the overall integrative analysis. J.R.D., V.T.L., C.A., F.A. and J.X. led Hi-C analysis. J.X., S.F., D.V.B., Y.W., R.C., J.B. and F.Y. led optical mapping and WGS analysis. V.D., T.S., J.C. and D.G. led replication timing analysis. Y.Z., H.O., B.L., and J.R.D. performed Hi-C experiments. D.P., S.H., C.E., and D.O. performed Hi-C on TH1 cells. G.Y., L.Z., H.Y., T.L., S.I., L.A., C.P., R.K., M.B., K.L., M.D., J.S., D.G. analyzed data. J.R.D., J.X., V.D., F.S., F.A., R.H., W.S.N., J.D., D.G., and F.Y. wrote the manuscript.

## Data Access

Hi-C and replication timing data generated in this study have been deposited to ENCODE portal (http://encodeproiect.org/). Details can be found in the supplementary method section. WGS data has been deposited to SRA (access code PRJNA380394). Replication time data can be visualized at http://replicationdomain.com. Hi-C data can be visualized at http://3dgenome.org.

